# Development and Evaluation of a Colloidal Gold Immunofiltration Assay for Penside Diagnosis of Visceral Schistosomosis in Cattle

**DOI:** 10.1101/2022.05.12.491591

**Authors:** Shivani Mamane, N. Jeyathilakan, Bhaskaran. Ravi. Latha, T. M. A. Senthilkumar

## Abstract

**Background:** Visceral schistosomosis caused by *Schistosoma spindale* is an economically important chronic wasting snail-borne trematode infection which often goes undiagnosed in field condition due to lack of efficient diagnostic assays. ELISA and Colloidal gold immunofiltration assay were used to evaluate the state of active visceral schistosomosis in cattle through detection of antibodies against *S. spindale* ES antigen.

**Methodology/Principle Findings:** The present study included development and evaluation of enzyme linked immunosorbent assay (ELISA) and colloidal gold immunofiltration assay (CGIFA) with conventionally prepared *S. spindale* excretory-secretory (ES) antigen using postmortem mesenteric examination as a reference standard. Statistical analysis was performed using Cochran Q test, Chi-square test and kappa statistics which showed no significant difference between the developed tests and strong agreement with the postmortem examination which was considered as gold standard test. CGIFA showed sensitivity, specificity and accuracy of 92.98 %, 96.74 % and 95.55 % respectively whereas that of ELISA was 94.73 %, 95.12 % and 95% respectively.

**Conclusion/Significance:** CGIFA was found to be simpler, rapid, effective, less expensive and required least expertise for interpretation of results. It can be used as portable diagnostic device for rapid diagnosis of *S. spindale* infection at field condition making it an ideal pen side diagnostic kit.

## Introduction

Schistosomosis is a snail-borne trematode infection which is identified as one of the major economically important diseases of human beings, domestic and wild animals mainly present in tropical and subtropical regions of the world (1). Animal schistosomosis is widespread throughout Asia and Africa. It has been conjectured that around 530 million cattle live in endemic areas and about 165 million cattle are infected with schistosomes globally (2). It is now well recognized as the fifth major helminthosis of domestic animals in the Indian subcontinent (3).

Visceral schistosomosis caused by *S. spindale is* a neglected chronic wasting blood fluke illness of livestock wherein the adult worms are obligate parasite of blood vascular system. This chronic disease is accompanied with symptoms such as frequent diarrhoea with traces of blood and mucous, anemia, edema, substantial reduction in productivity and emaciation which are insufficient clinical signs to distinguish the illness from other debilitating diseases (4). It causes severe economic losses due to liver condemnation and produces a high morbidity (5). It has been reported in India, Sri Lanka, Indonesia, Malaysia, Thailand, Laos and Vietnam. It resides in the mesenteric veins of cattle, buffalo, sheep and goats causing visceral or hepatointestinal schistosomosis. Abattoir surveys revealed that 30-68 per cent of bovines in South Indian states such as Karnataka, Tamil Nadu, Kerala and Telangana are infected with visceral schistosomosis (3, 6-8). *Schistosoma spindale* is one among the mammalian schistosomes responsible for cercarial dermatitis in human beings of India (9). Cercarial dermatitis outbreaks have been reported in India mainly from Chennai (erstwhile Madras), Assam, Madhya Pradesh and Central India (9-12).

Microscopic examination of faeces/rectal pinch for ova or miracidium is generally considered for diagnosis, mainly because of high specificity. However faecal egg detection method is considered unreliable due to low egg number output (13). These parasitological tests are also time consuming and limited in sensitivity in subclinical cases and at times the ova are trapped in the mucous of faecal matter and goes unnoticed in acute cases. Because of poor efficacy and dependability of the coprology based detection, the prevalence of this economically important disease is underestimated (14) and accordingly the situation calls for development of immunodiagnostic assays in order to supplement or replace the existing less sensitive faecal examination techniques. Currently for diagnosis of schistosomosis immunological tests such as enzyme linked immunosorbent assay, counter current immunoelectrophoresis, indirect haemagglutination, immunofluorescence assay are used which are used with certain shortcomings such as requirement of sophisticated laboratory equipment, reagents and trained personnel in addition to the time consuming protocols.

In contrast to human schistosomosis which is Neglected Tropical Disease (NTD), very few immunological studies have been conducted in animal schistosomosis (15). There is a necessity to prioritize the control of visceral schistosomosis in cattle since studies have proved that hepatic schistosomosis is more widely prevalent than nasal schistosomosis or fasciolosis and is often underdiagnosed (13).

The present investigations was aimed at developing and evaluating the CGIFA and comparing the results with ELISA so as to develop a simple, sensitive penside diagnostic test, CGIFA which could lead to improved rapid and accurate diagnosis using ES antigen for the sero-detection of antibodies evoked by *S. spindale* infection in cattle.

## Materials and Methods

### ES antigen preparation

*Schistosoma spindale* worms were collected by screening the mesentery samples obtained from Perambur slaughter house, Chennai, Tamil Nadu, India (13.1038° North, 80.2612° East). The worms were checked for viability under the microscope and mixed adult *S. spindale* worms were soaked in Phosphate Buffer Saline for two hours at room temperature followed by overnight incubation at 4°C. After incubation under sterile condition, the worms were carefully removed, the remaining filtrate was collected and centrifuged at 10,000 xg for 30 minutes at 4°C and the supernatant was used as ES antigen (16, 17). The protein concentration of ES antigen was estimated using protein estimation kit by Bradford method (Merck GeNei™, Bangalore).

### Collection of serum samples

A total of 180 serum samples were collected from the cattle at Perambur slaughter house, of which 57 were found to be positive for the *S. spindale* infection by mesenteric examination while other 123 were from the parasite free cattle.

Hyper immune serum was raised against ES antigen of *S. spindale* in two adult New Zealand white rabbit (3-4 kg) by repeated immunizations and used for standardization of CGIFA (18).

### Ethical statement

Permission was obtained from Ethical committee of Madras Veterinary College, Chennai bearing the Proposal No. 82/SA/IAEC/2019 dated 21-11-2020.

### Enzyme linked immunosorbent assay (ELISA)

ELISA was used for the detection of serum antibodies against *S. spindale*. The standardized ELISA was evaluated using 200 ng/well ES antigen, 1:50 dilution of test sera sample and 1:5000 dilution of HRP conjugate. The mean OD values of ELISA performed with known negative samples was calculated and cut-off value for determining positive and negative samples was obtained by using formula, mean OD plus 3 times standard deviation.

### Colloidal gold immunofiltration assay (CGIFA)

CGIFA was carried out by considering the one developed for the diagnosis of human schistosomosis (19). Nitrocellulose (NC) membrane of 0.45µm size was soaked in 80 per cent alcohol for 10 minutes. The membrane was air dried and cut into 3.5 × 2.5 cm size pieces. The NC membrane pieces were pressed tightly on top of the water absorbing pad (Advanced Microdevices Pvt Ltd, Ambala Cantt, India). 5 µl of ES antigen (80µg/ml) was added to NC membrane as small dot using flow directing funnel in the device. Antigen coated flow through device were taken to standardize the test. The NC membrane of flow through device was added with 100 µl washing buffer (PBS, 0.5 % BSA, 0.05 % Tween-20, 0.2 % Sodium azide) through flow directing funnel for a minute. Then, 20 µl volume of hyper immune serum was added onto the device and allowed to absorb completely. After that 50 µl of 1:5 dilution Protein A colloidal gold conjugate (Sigma, USA), which is used as an indicator was added to the test hole and was allowed to soak for 2 minutes. Finally 100 µl of washing buffer was added to remove the unbound conjugate. The whole procedure was completed within 10-15 minutes and the results were determined with unaided eyes. Likewise, negative control test using known negative serum was done to compare the results.

### Comparison of CGIFA and ELISA

Samples tested by ELISA were tested using the immunochromatographic assay.The known positive and negative cattle serum samples tested by ELISA were subjected to the developed immunochromatographic assay to know the respective sensitivity, specificity and accuracy of the tests using the gold standard test.

### Statistical analysis

A contingency 2×2 table was created for the assessment of sensitivity, specificity, efficiency and predictive values of ELISA and CGIFA for the diagnosis of visceral schistosomosis caused by *S. spindale* using 180 serum samples of cattle by keeping postmortem examination as a gold standard.

The results of parameters between each test were compared with gold standard and analyzed by the Cochran Q test, Chi-square test and Kappa statistics using SPSS software (Version SPSS 26). Cochran Q test was applied for comparison of the response of the two immunodiagnostic assays with gold standard as samples were related; Kappa statistics and Chi-Square tests were calculated to know the degree of agreement between the developed tests.

## Results

Out of 180 mesentery samples, 57 were positive during postmortem examination showing overall 31.67 per cent infection rate. Serodiagnosis of visceral schistosomosis caused by *S. spindale* was done by standardization of ELISA using known positive and negative bovine sera whereas CGIFA was standardized using negative rabbit sera and hyper immune sera raised in rabbits. Among 180 cattle serum samples collected, 57 were found positive and 123 were negative by postmortem examination of mesentery which was considered as gold standard for evaluating ELISA and CGIFA.

### ELISA

Out of 57 positive and 123 negative samples, 54 were positive and 117 were negative by ELISA respectively. The sensitivity, specificity, positive predictive value, negative predictive value, accuracy, concordance, false positive rate, false negative rate, kappa value and Youden index of ELISA in detecting serum antibodies in cattle was 94.73 per cent, 95.12 per cent, 90 per cent, 97.5 per cent, 95 per cent, 94.48 per cent, 4.88 per cent, 5.27 per cent, 0.8743 and 0.89 respectively.

**Table 1:** Evaluation of Enzyme linked immunosorbent assay (ELISA) for detection of *Schistosoma spindale* infection in cattle (n=180)

|  | Mesentery examination |  |  |
| --- | --- | --- | --- |
| ELISA | + | - | Total |
| + | 54 | 6 | 60 |
| - | 3 | 117 | 120 |
| Total | 57 | 123 | 180 |

Chi-square value=138.44\*\*, $K = 0.8743$

### CGIFA

Out of 57 positive and 123 negative samples, 53 were found positive and 119 were negative by CGIFA respectively. The sensitivity, specificity, positive predictive value, negative predictive value, accuracy, concordance, false positive rate, false negative rate, kappa value and Youden index of FTT in detecting serum antibodies in cattle was 92.98 per cent, 96.74 per cent, 92.98 per cent, 96.74 per cent, 95.55 per cent, 95.56 per cent, 3.26 per cent, 7.02 per cent, 0.8973 and 0.892 respectively.

**Figure 1.**
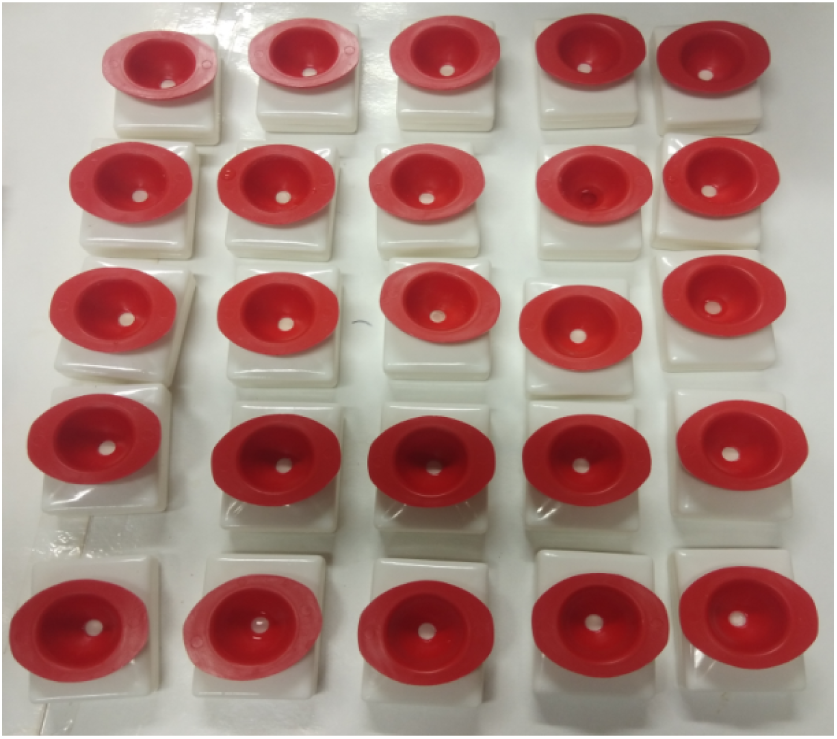
Evaluation of CGIFA.

**Figure 2.**
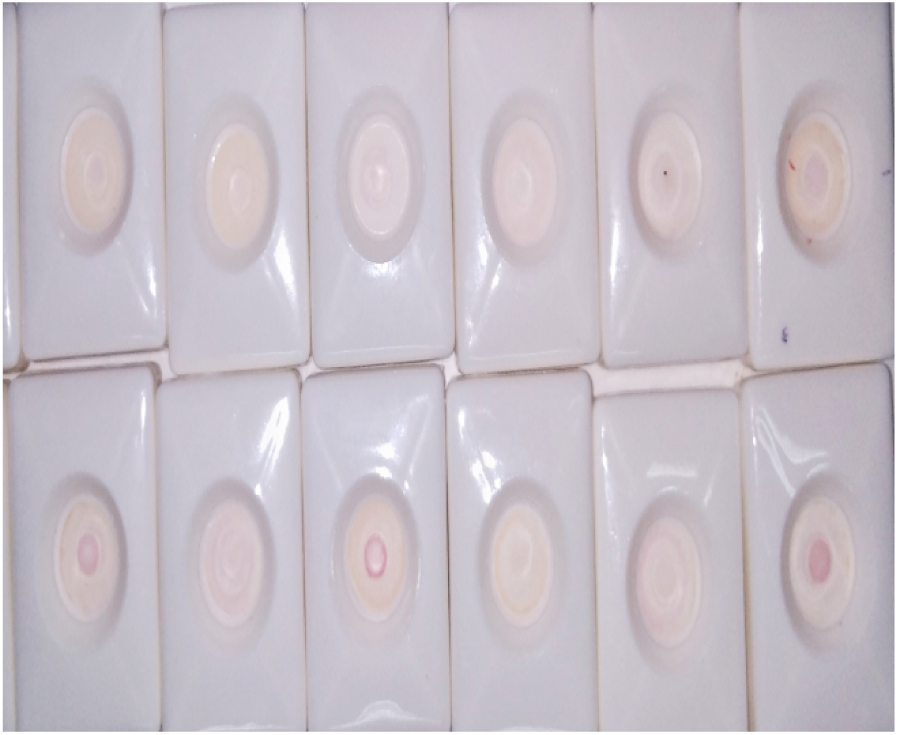
Development of colour in positive infection of the developed CGIFA.

**Table 2:** Evaluation of Colloidal gold immunofiltration assay(CGIFA) for detection of *Schistosoma spindale* infection in cattle (n=180)

|  | Mesentery examination |  |  |
| --- | --- | --- | --- |
|  | + | - | Total |
| CGIFA |  |  |  |
| + | 53 | 4 | 57 |
| - | 4 | 119 | 123 |
| Total | 57 | 123 | 180 |

Chi-square value=144.93\*\*, $K=0.8973$

Comparative evaluation of the developed tests unveiled that CGIFA showed higher specificity and accuracy than ELISA whereas ELISA showed higher sensitivity than CGIFA. Cochran Q test and Chi-Square tests revealed no significant difference between the developed serodiagnostic assays. Kappa statistics also showed the presence of strong agreement between the tests when compared with the gold standard.

## Discussion

It is clearly evident that though the clinical cases of visceral schistosomosis are minimal and subclinical cases do exists. It is difficult to detect this condition since affected animal shows no obvious sign of disease and continues to be a carrier, spreading the disease (3). Considering the schistosome species, finite amount of work has been conducted on immunodiagnosis of *S. spindale* in cattle by using whole worm antigen, soluble egg antigen and ES antigen for detection of serum antibody or antigen with varying sensitivity and specificity (19-22).

The excretory secretory, tegumental and gut proteins of adult schistosomes have been reported to be a rich source of potential immunogens (23). Schistosome ES proteins have been shown to play an important role in modulating host immune responses (16). The present study involved preparation of ES antigen against the whole worm homogenates because ES antigens evoke a considerably higher degree of protective immunity in lab animals than the somatic antigen derived from whole worm (24). Lack of cross reactivity and the diagnostic specificity of ES antigen was confirmed and strong recommendation for the incorporation of ES antigen in various immunodiagnostic assays for detection of *S. spindale* was advised (17).

ELISA was performed for studying the seroprevalence of *S. spindale* infection in different agro climatic zones of South India, using 300 ng ES antigen concentration per well, 1:200 dilution of the serum samples and 1:2000 dilution of HRP conjugate and an overall sensitivity and specificity of 96 and 100 per cent was observed (22). The study on efficacy of Dot-ELISA using whole worm antigen, ES antigen and soluble egg antigen of *S. spindale* at various concentration of 100 ng, 250 ng and 100 ng respectively showed cent per cent sensitivity and specificity (25). However, in another study evaluation of Dot-ELISA carried out using ES antigen of *S. spindale* showed 80.48 and 78.57 per cent sensitivity and specificity respectively (21). Evaluation of Sandwich ELISA using whole worm antigen of *S. spindale* for faecal antigen detection revealed sensitivity and specificity of 88 and 90 per cent (19). The variation in the results could be due to the differences in methodology used, concentrations used for checker board analysis, interpretation of the results and the nature of antigenicity of proteins used in the assays. However, conventional ELISA has some drawbacks including the need of trained personals, requirement of specific and costly equipment, lack of reproducibility due to plate to plate variation and limitation in field level application.

Various kind of immunofiltration assays have been developed for diagnosis of human schistosomosis caused by *S. japonicum* using soluble egg antigen conjugated with the red colloidal dye and blue colloidal dye (18, 26). The sensitivity and specificity was found to be 94.1 and 96.7 per cent whereas 98 and 99.4 per cent using the respective dyes in their developed assay. Dot-Dye immune assay was described for detection of *S. mansoni* with antigen conjugated with textile dye, this assay showed 93.4 per cent sensitivity and 100 per cent specificity showing no significant difference when compared with ELISA and Dot-ELISA (27). The dipstick with latex immunochromatographic assay to detect *S. japonicum* was devised with 95.1 and 94.91 per cent of sensitivity and specificity respectively (28), when compared with ELISA no significant difference was found and this observation goes hand in hand with that of the present investigation. The variation in the sensitivity and specificity mainly depends on the gold standard test considered, species of Schistosome and antigen involved in the assay, conjugate colloidal employed and its ability to bind with the test antibodies, test procedure. The available literature indicated that no study has so far been attempted on serodiagnosis of visceral schistosomosis in cattle using ES antigen based CGIFA. Hence, this study is probably the original attempt and first of its kind in the detection of *S. spindale* infection in cattle. Use of the dyes instead of Protein A colloidal gold can be employed to further cut down the cost of the test and for future application in large scale screening in field conditions.

Among the two assays, CGIFA is considered as the simple test with the advantages such as minimum amount of antigen required for testing a large amount of samples. Statistically no significant difference was found between the ELISA and FTT in the detection of antibodies against *S. spindale* infection (*P*>0.05). CGIFA involves less expensive equipment without involving any incubation process and can be done at room temperature within 10 minutes. It was found to be less complex, rapid, effective, less expensive, easy to apply and needs less expertise for result interpretation. The evaluated assays can be used for large scale screening of sera for surveillance studies of visceral schistosomosis caused by *S. spindale* in cattle. CGIFA can be used as portable diagnostic device for rapid diagnosis of *S. spindale* infection at field condition as a pen side diagnostic kit.

## Notes

### Competing Interest Statement

The authors have declared no competing interest.

